# Xeno-free induced pluripotent stem cell-derived neural progenitor cells for *in vivo* applications

**DOI:** 10.1101/2022.01.18.476253

**Authors:** Ruslan Rust, Rebecca Z. Weber, Melanie Generali, Debora Kehl, Chantal Bodenmann, Daniela Uhr, Debora Wanner, Kathrin J. Zürcher, Hirohide Saito, Simon P. Hoerstrup, Roger M. Nitsch, Christian Tackenberg

## Abstract

Cell-based therapies are a promising treatment paradigm for neurodegenerative diseases and other brain injuries. Despite recent advances in stem cell technology, major concerns have been raised regarding the feasibility and safety of cell therapies for clinical applications. Here, we generate good manufacturing practice (GMP)-compatible neural progenitor cells (NPCs) from transgene- and xeno-free induced pluripotent stem cells (iPSCs) that can be smoothly adapted for clinical applications. The produced NPCs have a stable gene-expression over at least 15 passages and can be scaled for up to 10^18^ cells per initially seeded 10^6^ cells. To ensure a pure NPC population for *in vivo* applications, we reduce risks of iPSC contamination using micro RNA-switch technology as a safety checkpoint. Using lentiviral transduction with a fluorescent and bioluminescent dual-reporter construct, combined with non-invasive *in vivo* bioluminescent imaging, we longitudinally tracked the grafted cells in healthy wild-type and genetically immunosuppressed mice as well as in a mouse model of ischemic stroke. Long term in-depth characterization revealed that transplanted cells have the capability to survive and spontaneously differentiate into functional and mature neurons throughout a time course of a month.

## Introduction

Neurodegenerative diseases and other brain injuries represent a huge personal and economic burden [1]. Cell-based therapy is considered an emerging treatment paradigm which has the potential to regenerate damaged tissue and restore brain function. While safety and feasibility of administering different types of stem cell therapies seem to be reasonably proven, its efficacy remains uncertain due to conflicting results in clinical trials [2]. A major limitation in a clinical set-up is the choice of a reliable and scalable cell source as dosing of up to 1 billion cells may be required for a single transplantation [3]. Most clinical trials rely on the use of primary mesenchymal stem cells (MSCs) due to their ubiquitous presence in the adult body. However, MSCs have major drawbacks with regard to heterogeneity, scalability and differentiation potential, especially into cell types of the neural lineage. Recent developments in induced pluripotent stem cell (iPSC) technology facilitated the infinite generation of neural progenitor cells (NPCs), a more homogenous cell source for brain regeneration through cell replacement, neuroprotection and/or immunomodulation [4].

While advancing the iPSC technology towards clinical reality, challenges have emerged regarding the presence of transgenes following reprogramming, xenogenic factors in culture media and the risk of teratoma formation from undifferentiated cells in graft recipients [5].

The use of the non-integrative RNA Sendai virus vector enabled reprogramming free of vector and transgene sequences [6]. Moreover, improved protocols for NPC differentiation and culture enabled propagation in defined media omitting xenogenic factors [7]. The risk of teratoma formation can be reduced by applying highly efficient differentiation protocols and/or effective purification techniques, such as the RNA Switch technology to specifically eliminate pluripotent residuals from cultures [8,9]. This technique is based on the presence of cell-specific micro RNA (miRNA), that can be used to either enrich or eliminate a specific cell type from a cell culture. These advancements increased the safety and feasibility of NPCs for preclinical and clinical *in vivo* applications. However, there is still poor understanding of the spatiotemporal kinetics, survival capabilities and differentiation phenotype of transplanted cells in an *in vivo* system that need to be answered before translating cell therapies into standard clinical applications.

Here, we generate and characterize NPCs from iPSCs without genomic modification and under a xeno-free, chemically defined environment. The generated NPCs show a stable gene expression over 15 passages and capability to spontaneously differentiate into functionally active neurons *in vitro*. We further ensured a pure NPCs population by eliminating residual iPSCs using RNA-switch technology. To characterize the fate of NPCs *in vivo*, cells were transduced using a dual-reporter system consisting of bioluminescence and fluorescence reporters. Transplanted grafts were longitudinally tracked and phenotyped one month after transplantation. The differentiation profile of NPCs revealed a strong preference to the neuronal fate with the expression of various neuronal markers. On the basis of the analysis in this study, we demonstrate the reliable generation of highly pure NPCs with a reduced risk of pluripotent residuals for long-term *in vivo* applications in the mouse.

## Materials and Methods

### Generation and maintenance of NPCs

NPCs were generated as previously described [10] with modifications. In detail, 80.000 iPSCs per well were plated on day −2 on a 12-well plate, coated with Vitronectin (StemMACS iPS-Brew XF (Miltenyi), supplemented with 2uM Thiazovivin) and incubated overnight. On day −1, cells were washed once in PBS and the medium was changed to fresh StemMACS iPS-Brew XF. On day 0, NPC differentiation was induced by washing cells in PBS and changing medium to Neural Induction Medium 1 (50% DMEM/F12, 50% Neurobasal medium, 1x N2-supplement, 1x B27-supplement, 1x Glutamax, 10 ng/ml hLIF, 4 μM CHIR99021, 3 μM SB431542, 2 μM Dorsomorphin, 0.1 μM Compound E). On day 1 fresh medium was added. On day 2 cells were washed with PBS and the medium was changed to Neural Induction Medium 2 (50% DMEM/F12, 50% Neurobasal medium, 1x N2-supplement, 1x B27-supplement, 1x Glutamax, 10 ng/ml hLIF, 4 μM CHIR99021, 3 μM SB431542, 0.1 μM Compound E) followed by medium change every day for 4 days. On day 6 cells were split onto pLO/L521-coated plates in Neural Stem cell Maintenance Medium (NSMM, 50% DMEM/F12, 50% Neurobasal medium, 1x N2-supplement, 1x B27-supplement, 1x Glutamax, 10 ng/ml hLIF, 4 μM CHIR99021, 3 μM SB431542). NSMM medium was changed every day and cells were split when reaching 80-100% confluency. For the first 6 passages 2μM Thiazovivin was added after splitting. From passage 2 on NSMM medium was supplemented with 5 ng/ml FGF2. Find details about media and cell culture materials in Suppl. Tables 1-4.

### Neural differentiation of NPCs

NPCs were seeded 200’000 cells/ml in NSMM + 5 ng/ml FGF2 on 24-well plates. Differentiation was induced by withdrawal of small molecules CHIR99021, SB431542, hLIF and FGF2. Cells were cultured for 21 days in NSMM without small molecules, hLIF and FGF2 with daily media changes during the first week and media changes every other day or biweekly in the second or third week, respectively.

### NPC transduction for in vivo tracking

For in vivo cell tracking, dual-reporter lentiviral plasmids pLL410_EF1a-rFLuc-T2A-GFP-mPGK-Puro (LL410PA-1) and pLL-CMV-rFLuc-T2A-GFP-mPGK-Puro (LL310PA-1) were obtained from System Bioscience. Lentiviral generation was carried out as previously described [11]. For NPC transduction, 650’000 cells/well were seeded in NSMM + 5 ng/ml FGF2. After 24 h, 20 μl virus per well was added during medium change (=1/100 virus per well).

### NPC transduction and calcium imaging

Adeno-associated viruses (AAV) at titer ≥ 1×10^13^ vg/ml, expressing mRuby2 and GCaMP6s under the human synapsin promoter (pAAV-hSyn1-mRuby2-GSG-P2A-GCaMP6s-WPRE-pA) was purchased from Addgene (50942-AAV1). NPCs were differentiated to neural cells as described above and transduced by adding 1.44 μl virus solution per well in a μ-Slide 4 Well 5 days after cell seeding. For neuronal stimulation KCl was added during imaging to a final concentration of 60 mM. Images were acquired using Leica SP8 confocal microscope, equiped with 10x HC PL FLUOTAR 0.30 DRY objective. Image settings for GCaMP6s were the following: 488 nm laser excitation (Argon laser); xyt scan mode; 512×512 pixel; 15min time-lapse imaging with 1.29s intervals; 200Hz. Image settings for mRuby were the following: 561 nm laser excitation (DPSS); xyz scan mode; 1024×1024 pixel; 100Hz.

### Image quantification for mRuby and GCamp6

For mRuby quantification, fluorescence signal intensity of mRuby was measured in randomly selected regions of interest (ROIs) and normalized to the DAPI cell count using Fiji (ImageJ). For GCamp6 quantification: at least 20 differentiated neural cells per condition were identified and outlined as ROIs using Fiji (ImageJ). Fluorescence signal intensity in these ROIs was continuously measured over 500 seconds.

### qPCR

RNA extraction was carried out using Quiagen RNeasy kit according to the manufacturer’s recommendations. qPCR was performed using SYBR green kit (iTaq Universal SYBR Green Supermix from Biorad) containing 0.5 μM of each primer with the following cycling conditions (hold stage: 95°C, 10 min, 1 cycle; PCR stage (95°C, 15s, 60°C 1 min; 95°C 15s, 40 cycles; Melting curve (95°C, 15s, 60°C, 1 min). Human total brain Total RNA (ThermoFisher, QS0611) served as a positive control.

### Immunocytochemistry

ICC was performed with 4% paraformaldehyde (PFA) /4 % sucrose fixed cells, whereby donkey serum was used for blocking and staining solution. Incubation of primary antibody namely NPC specific markers Pax6, Sox1 and Nestin as well as iPSC specific marker Oct4, was performed at 4 °C overnight. The next day the secondary antibody was incubated for 2 h at room temperature (RT) followed by DAPI staining. Confocal microscopy was used for imaging.

### Flow cytometry

Cells were thawed and incubated in FACS buffer (0.1% BSA in PBS) with live/dead Fixable Near-IR staining (Molecular probes) for 30 min at 4°C. Subsequently, cells were fixed in 1 % PFA for 20 min at RT and permeabilized with FACS buffer containing 0.5% Saponin for 20 min at 4 °C. Intracellular antibodies namely Pax6, Nestin, Sox1 and Oct4 were then used for staining for 30 min at 4 °C. Cells were acquired using a LSR Fortessa (BD Bioscience) and the data were analyzed by FlowJo software (Tree Star)

### RNA switch

The RNA switch was carried out as previously described [8]. Briefly, template DNA for *in vitro* transcription of normal mRNA was amplified by PCR from a vector encoding for either Barnase, Barstar, or puromycin using appropriate primers with the T7 promoter and poly(A) tail. The template of miRNA-responsive OFF switch (Barstar) was amplified from the same vector by PCR using the primer containing miRNA anti-sense sequence at 5’UTR. For the miRNA-responsive ON switch (Barnase), the open reading frame, poly(A) tail, and miRNA anti-sense sequence were amplified from the same vector by first PCR. Then, the PCR product and extra sequence were fused in this order by a second PCR. All template DNAs were purified using the MinElute PCR Purification Kit. The RNAs were transcribed for 6 hours at 37C using MEGAScript T7 Transcription Kit using 1-Methylpseudouridine-5’-Triphosphate and Anti Reverse Cap Analog, ARCA. The transcribed RNA was treated with Turbo DNase I and antarctic phosphatase to remove template DNA and purified by RNeasy MinElute Cleanup Kit (QIAGEN). The concentration was determined by Quibit microRNA Assay Kit. A day before transfection, cells were passaged as detailed above and seeded onto PLO/L521 coated plates. The following day, miRNA was transfected using Lipofectamin RNAiMAX Transfection Reagent. The transfection complex was added to the cells in a drop-wise manner and the plate was agitated before being placed into the incubator, followed by a medium change after 4h. To remove non-transfected cells, a puromycin selection was carried out by treatment with 2 μg/ml puromycin for 24h. Cells were finally analysed by flow cytometry.

### In vitro luciferase assay

Cells were seeded on 24-well plate and a serial diluted luciferin standard curve (starting concentration was 300 ug/ml) was made. After incubating cells for 10 minutes at 37 °C bioluminescence was measured with Tecan M1000 pro.

### NPC viability analysis

To determine viability of NPC after thawing and before transplantation, NPCs (150000 cells / ml in PBS) were stored on ice and cell viability was measured using the Vi-Cell XR Cell Viability Analyzer.

### Animals

All animal experiments were performed in accordance with governmental and institutional (University of Zurich) guidelines and had been approved by the Cantonal Veterinary Office of Zurich (License number: 31687). Immunocompromised recombination activating gene 2 knockout mice (Rag2^-/-^), non-obese diabetic SCID gamma mice (NSG) and wild type (wt) mice with a C57BL/6 background were used (10 – 14 weeks old, female and male). Animals were housed in standard Type II/III cages at least in pairs in a temperature and humidity-controlled room with a constant 12/12h light/dark cycle (light on from 06.00 a.m. until 6:00 p.m.) and food/water ad libitum.

### Cell transplantation

Cells were transplanted as previously described [12]. In brief, NPCs at passage number ≥ 11 were used in all experiments. At time of cell transplantation, GFP^+^/Luc^+^ NPCs were diluted with a final concentration of 8 × 10^4^ cells/μL in sterile 1x PBS (pH 7.4, without calcium or magnesium; Thermo Fisher Scientific) and stored on ice until transplantation. Mice were anesthetized using isoflurane (4% induction, 1.5% maintenance; Attane, Provet AG). Analgesic (Rimadyl; Pfizer) was administered subcutaneously prior to surgery (5 mg/kg body weight). Animals were placed in a stereotactic frame (David Kopf Instruments), the surgical area was sanitized, and the skull was exposed through a midline skin incision to reveal *lambda* and *bregma* points. Coordinates were calculated (the coordinates of interest chosen for this protocol were: AP: + 0.5, ML: + 1.5, DV: −0.8 relative to bregma) and a hole was drilled using a surgical dental drill (Foredom, Bethel CT). Mice were injected with 1.6 × 10^5^ cells (2μL/injection) using a 10-μL Hamilton microsyringe (33-gauge) and a micropump system with a flow rate of 0.3 μL/min (injection) and 1.5 μl/min (withdrawal). In addition, serial dilutions (1.8 × 10^5^, 1 × 10^5^, 6 × 10^4^, 1.2 × 10^4^, 6 × 10^3^) of NPCs were injected into the cortex (AP: + 0.5, ML: + 1.5, DV: −0.8 relative to bregma) to determine the minimum cell number that can be detected *in vivo*. Wounds were sealed using a 6/0 silk suture and mice were allowed to recover in cages with heating pads. For postoperative care, all animals received analgesics (Rimadyl, Pfizer) for at least 2 days after surgery.

### Photothrombotic stroke

The induction of a photothrombotic stroke was carried out as previously described in wild-type (wt) mice [12–16] and in NSG and Rag2^-/-^ mice. In brief, mice were anesthetized using isoflurane (4% induction, 1.5% maintenance, Attane, Provet AG). They received analgesic (Novalgin, Sanofi) 24 h prior to the start of the procedure, administered via drinking water. A photothrombotic lesion was induced in the right sensorimotor cortex. Briefly, animals were fixed in a stereotactic frame (David Kopf Instruments) and the surgical area (from the back of the neck up to the eyes) was shaved and sanitized (Betadine, Braun). Anaesthesia was maintained using a face mask. Eye lubricant (Vitamin A, Bausch&Lomb) was applied, and body temperature was constantly kept between 36° and 37° C using a heating pad. The skull was exposed through a midline skin incision. A cold light source (Olympus KL 1,500LCS, 150W, 3,000K) was positioned over the right forebrain cortex (anterior/posterior: −1.5– +1.5 mm and medial/lateral 0 mm to +2 mm relative to Bregma). 5 minutes prior to illumination, Rose Bengal (15 mg/ml, in 0.9% NaCl, Sigma) was injected intraperitoneally. Subsequently, the exposed area was illuminated through the intact skull using an opaque template with an opening of 3×4mm. After 10.5 minutes, light exposure was stopped and the wound was closed using a 6/0 silk suture. For postoperative care, all animals received analgesics (Novalgin, Sanofi and Rimadyl, Zoetis) for at least 3 days after surgery.

### Immunohistochemistry

For histological analysis, animals were euthanized by intraperitoneal application of pentobarbital (150mg/kg body weight, Streuli Pharma AG) and transcardially perfused with Ringer solution (containing 5 ml/l Heparin, B. Braun) followed by PFA (4%, in 0.2M phosphate buffer, pH 7). Brains were postfixed for approximately 4 h in the same fixative, then transferred to 30% sucrose for cryoprotection and stored at 4°C. Coronal sections with a thickness of 40 μm were cut using a sliding microtome (Microm HM430, Leica), collected and stored as free-floating sections in cryoprotectant solution at −20°C until further processing.

Brain sections were washed with 0.1M phosphate buffer (PB) and incubated with blocking solution containing donkey serum (5%, Sigma) in PB for 60 min at room temperature. Sections were incubated overnight at 4°C with either mouse anti-mitochondria antibody (Thermo Fisher Scientific, 1:200, # MA5-12017); mouse anti-nuclei antibody (Merck, 1:100), mouse anti-STEM101 antibody (Takara Bio Inc., 1:100) to stain for transplanted human NPCs. To identify transplanted cells at different developmental stages, the following antibodies were used: Goat anti-human NANOG (R&D systems, 1:200), rabbit Oct-4A (Cell Signaling Technology, 1:200), mouse Anti-PAX6 monoclonal antibody (Thermo Fisher Scientific), mouse Anti-NeuN Antibody (Merck, #MAB377), rabbit Anti-Neurofilament 200 antibody (Merck, #N4142), rabbit Anti-BetaIII-Tubulin antibody (Abcam, #ab18207), mouse MAP2 monoclonal antibody (Thermo Fisher Scientific), rabbit Anti-S100b antibody (Thermo Fisher Scientific). The primary antibody incubation was followed by 2 h incubation at room temperature with corresponding fluorescent secondary antibodies (1:500, Thermo Fisher Scientific). Nuclei were counterstained with DAPI (1:2000 in 0.1 M PB, Sigma). Sections were mounted in 0.1M PB on Superfrost PlusTM microscope slides and embedded in Mowiol. All images were taken using a Leica SP8 laser scanning confocal microscope equipped with 10x, 20x and 40x objectives. Images were processed using Fiji (ImageJ) and Adobe Illustrator CC.

### In vivo bioluminescence imaging

Bioluminescence imaging (BLI) experiments were performed with the IVIS Spectrum CT (PerkinElmer) as described before [12]. For long-term experiments, animals were imaged in regular intervals starting 24h after transplantation for up to 35 days. Animals used in the study to determine the minimal number of detectable cells were imaged only *once* 1.5h after transplantation. 30mg/ml D-luciferin potassium salt (PerkinElmer) was dissolved in NaCl (0.9%, B. Braun) and sterilized using a 0.22 μm syringe filter. Luciferin was injected intraperitoneally to each animal with a final dose of 300 mg/kg body weight before isoflurane anaesthesia (4% induction, 1.5% maintenance; Attane, Provet AG). Before the first BLI recording, animals were shaved on the head region. Image acquisition was performed for 20 min under the following setting: *Field of View: A, Subject height: 1.5 cm, Binning: 16, F/Stop: 2*. Exposure time (ranging from 1s to 60s) was set automatically by the system to reach the most sensitive setting. Imaging parameters and measurement procedures were kept consistent within and between subjects.

### In vivo imaging analysis

*In vivo* BLI data was analysed using the Living Image Software (LI 4.7.3) with size-constant ROIs as described before [12,17]. Briefly, the ROIs were manually drawn based on anatomical landmarks (eyes, ears and snout) on each image. Plotting and statistical analysis were performed using RStudio. The brain-specific signal was calculated and corrected for nonspecific signal taken from a ROI on the skin of the animal’s back and for noise taken from a ROI outside the mouse. Signal-to-noise ratio (SNR) was calculated by dividing the mean photon flux (ph/s/cm^2^/sr) by the standard deviation of the noise. Bioluminescent signal was calculated by subtracting the background flux from the mean photon flux.

### Statistical analysis

Statistical analysis was performed using RStudio 4.0. Sample sizes were designed with adequate power according to our previous studies. All data were tested for normal distribution by using the Shapiro-Wilk test. Normally distributed data were tested for differences with a two-tailed unpaired one-sample *t*-test to compare changes between two groups (GFP expression of transduced vs. control cells) as in Figure 4F. Multiple comparisons as in Figures 1C and Suppl. Figure 1A (NPC gene expression differences), Figures 2B,C,F (protein fluorescence intensity), Figure 3D (FACS iPSC signal) were initially tested for normal distribution with the Shapiro-Wilk test. The significance of mean differences between normally distributed multiple comparisons was assessed using Tukey’s HSD. Statistical significance was defined as \**p* < 0.05, \*\**p* < 0.01, and \*\*\**p* < 0.001.

**Figure 1:**
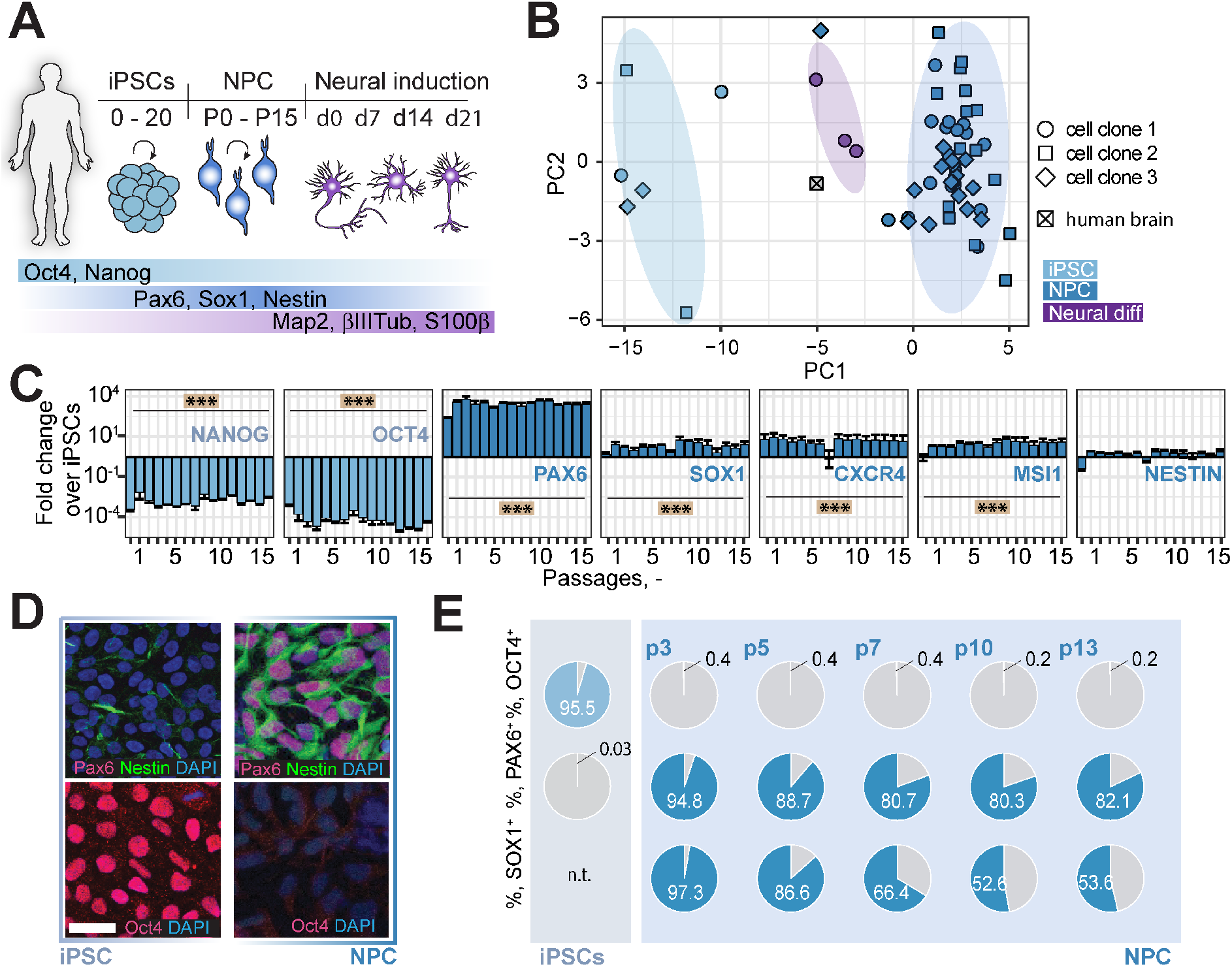
Generation and characterization of iPSC-derived NPCs. A: Schematic representation of the analyzed cell types, time frame and the main marker proteins used for characterization. B: Principal component analysis of qPCR data for NPCs (3 clones, dark blue) iPSCs (3 clones, light blue), NPCs 21d after neural differentiation (clone 1, violet) and human total brain RNA. Each symbol illustrates data from RNA extracted from one cell culture well. C: Gene expression of pluripotency marker (NANOG and OCT4) and NPC marker (PAX6, SOX1, CXCR4, MSI1, NESTIN) in NPCs over the course of 15 passages, measured by qPCR. D: iPSCs (left panel) and NPCs at passage 7-10 (right panel) stained for Pax6, Nestin, Oct4 and DAPI. E: Flow cytometry analysis of iPSCs (upper row) and NPCs at different passages (lower rows) for Oct4, Pax6 and Sox1. Pie charts illustrate percentage positivity (light/dark blue) for the respective marker and cell type. n.t.: not tested Neural diff: NPCs after neural differentiation; CXCR4: CXC-motif chemokine receptor-4; MSI1: Musashi-1; Scale bars: 50μm. Significance of mean differences between the groups was assessed using Tukey’s HSD. Asterisks indicate significance: *** P < 0.001.

## Results

### Generation of functional, pure iPSC-derived NPCs

To generate human neural cells with a high translational potential we differentiated human iPSCs into neural progenitor cells (NPCs) (Fig. 1A). Differentiation and maintenance of NPCs was carried out using dual-SMAD inhibition, also under xeno- and feeder-free conditions. NPCs were expanded and characterized over at least 15 passages. Gene expression analysis of iPSCs (3 different clones), NPCs (3 different clones) and differentiated cells at 7, 14 and 21 days after neural induction (1 clone) showed cell type-but not clone-specific clustering (Fig. 1B). Human total brain samples clustered next to neural cells which were differentiated from NPCs. Gene expression of pluripotency marker NANOG and OCT4 were highly downregulated in NPCs (all p<0.001), while marker for neural progenitors, i.e. PAX6, SOX1, CXCR4, and MSI1 were stably upregulated over 15 passages (all p <0.001, Fig. 1C). Immunostainings showed NPC positivity for Pax6 and Nestin but not for Oct4, while iPSCs were stained positive for Oct4 but not Pax6 (Fig. 1D). A mild staining was observed for Nestin These results were supported by flow cytometry analysis over several passages (Fig. 1E). The NPC differentiation protocol was further tested in two other iPSC clonal lines which gave similar results indicating robust NPC differentiation (Suppl. Fig. 1).

To determine the spontaneous neural differentiation potential of the generated NPCs, we withdrew CHIR99021, SB431542, hLIF and FGF2 from the NSMM over the course of 3 weeks. Immunocytochemical analysis of differentiated neural cells showed expression of neuronal markers Map2 and βIII-Tubulin, astrocyte marker S100β as well as NPC marker Pax6 but not pluripotency marker Oct4 (Fig. 2A upper row). The punctate staining of vesicular glutamate transporter (vGlut) and vesicular GABA transporter (vGat) along the neurites indicated the formation of synaptic contracts between neurons. Further, the number of βIII-Tubulin positive neurites (Fig. 2A, lower row) as well as the fluorescence intensity of Map2 and doublecortin (DCX, Fig. 2B) increase over time (all p < 0.001) suggesting a gradual increase in neural cell number and network complexity during differentiation. Compared to undifferentiated NPCs, differentiated cells showed a time-dependent increase in synaptic gene expression (vGAT: all p<0.001; vGLUT: p<0.05 (d14, d21)) (Fig. 2C). Astrocytic genes APOE and S100β were also partially upregulated (APOE: all p< 0.001, S100β: d14: p<0.001); and while the levels of oligodendrocyte marker slightly increased at d21 it did not reach significance (all p>0.05). Elevated levels of BRN2 and FOXG1 suggested the presence of excitatory cortical neurons. To confirm that differentiated neural cells were active and reacted to stimulation, we expressed mRuby2 and the calcium sensor GCaMP6s under the human synapsin 1 promoter using adeno-associated virus (AAV) (Fig 2D). Expression of mRuby was only observed in differentiated neural cells but not in undifferentiated NPCs (p<0.001, Fig. 2E, F) indicating activity of the synapsin 1 promotor only in differentiated cells. Neural cells showed higher GCaMP6s signals already at baseline and addition of 60 mM KCl immediately increased GCaMP6s fluorescence, analyzed by confocal live imaging (Fig. 2G, H).

**Figure 2:**
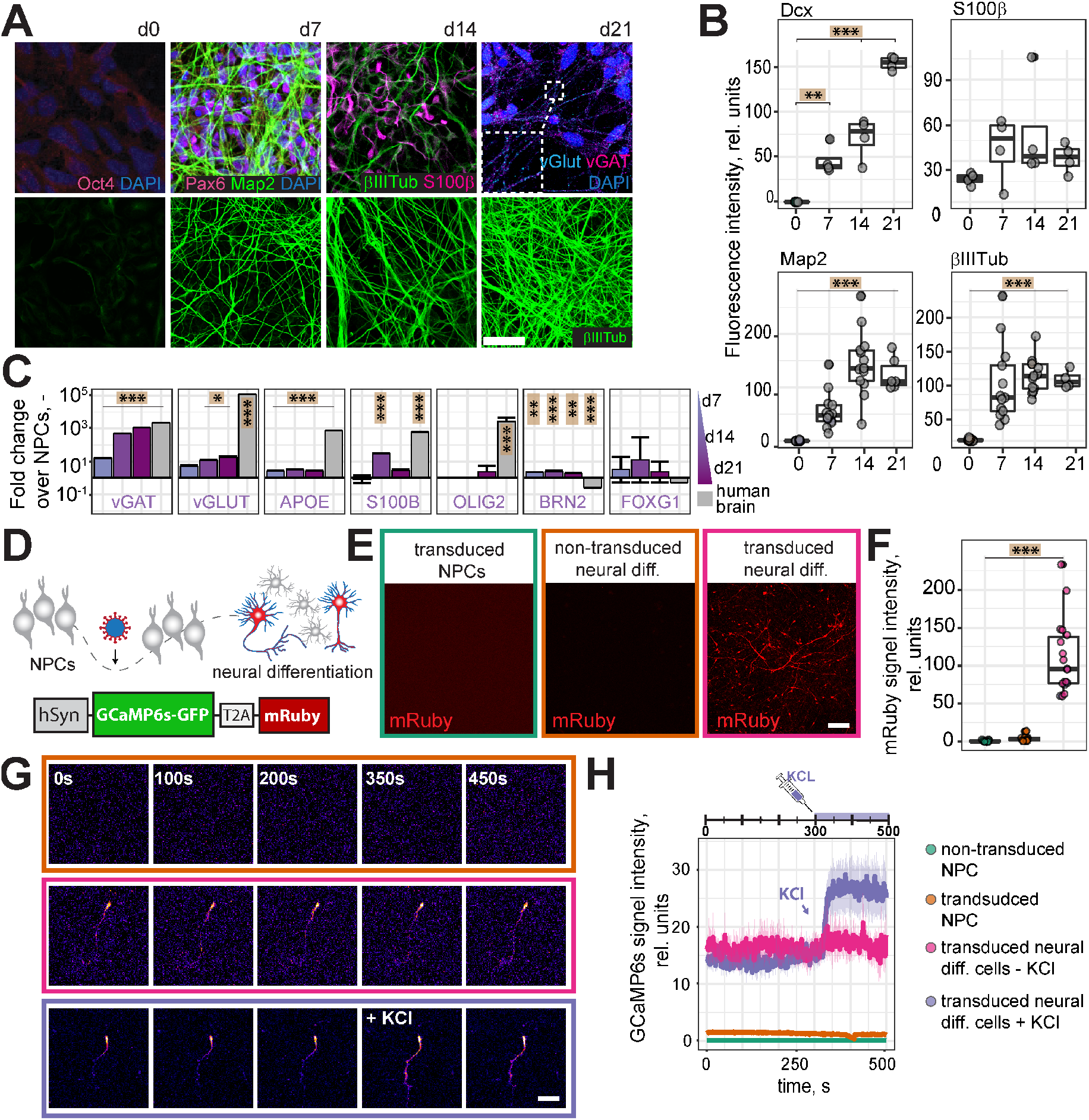
Neural differentiation of NPCs. A: Differentiated NPCs, at d21 after differentiation (upper row), stained for Oct4, Pax6, Map2, βIII-Tubulin, vGlut and vGat. Differentiated NPCs, at d0, d7, d14 and d21 after differentiation (lower row) stained for βIII-Tubulin. B: Quantification of fluorescence intensity of microscopy images of differentiated NPCs at d0, d7, d14 and d21 of differentiation, stained for Dcx, S100β, Map2 and βIII-Tubulin. C: Gene expression of neuronal (vGAT, vGLUT, BRN2, FOXG1), astrocyte (APOE, S100β) and oligodendrocyte marker (OLIG2) in differentiated NPCs at 7, 14 and 21d after neural differentiation, measured by qPCR. Total human brain RNA served as control. D: Experimental setup for determining the activity of differentiated NPCs and AAV construct for the expression of GCaMP6s and mRuby under the human synapsin promoter. E: Human synapsin promoter-based expression of mRuby in NPCs and differentiated neural cells. Confocal images of NPCs transduced with the AAV-mRuby construct (left), differentiated NPCs at d21 after differentiation (middle) and differentiated NPCs at d21 after differentiation and transduced with AAV-mRuby construct (right). F: Quantification of mRuby signal in transduced NPCs (left), non-transduced differentiated neural cells (middle) and transduced neural cells (right) from Fig.1G. G: Confocal time-lapse images of non-transduced, differentiated neural cells (upper row) and neural cells transduced with AAV-GCaMP6s construct (middle row). After 350s, cells were treated with 60 mM KCL (lower row). Scale bars: 50μm. H: Quantification of GCaMP6s signal intensity from confocal time-lapse images. vGAT: Vesicular GABA Transporter; vGLUT: vesicular glutamate transporter; Dcx: doublecortin; βIIITub: βIII-Tubulin; hSyn: human synapsin. Boxplots indicate the 25% to 75% quartiles of the data. Each dot in the plots represents one cell culture well and significance of mean differences between the groups was assessed using Tukey’s HSD. Line graphs are plotted as mean ± sem. Asterisks indicate significance: ** p< 0.01, *** p < 0.001.

Taken together, iPSC-derived NPCs generated under xeno-free conditions showed i) upregulation of typical NPC markers, ii) strong downregulation of pluripotency markers and iii) spontaneous differentiation into functional neurons and glial cells *in vitro*.

### iPSC-elimination using RNA switch

Remaining pluripotent cells are a major concern for iPSC-based cell therapies. To reduce the risk of iPSC residuals, we took advantage of the RNA switch technology to eliminate potential residual iPSCs. The RNA switch was designed to specifically detect miR-302a-5p that is highly expressed in hiPSCs and eliminate iPSCs in cell cultures [8,18]. Since our differentiation protocol already generated a highly pure population of NPCs, we tested the efficacy of the RNA switch by adding 10% and 20% of iPSCs to the NPC cultures (Fig 3A). Under both conditions we were able to reduce the iPSC-content of our cultures to <1% of Oct4-positive iPSCs, as detected by flow cytometry. A negative control RNA switch, in which cells were lipofected with no RNA, did not affect the amount of added iPSCs (Fig. 3 B, C).

**Figure 3:**
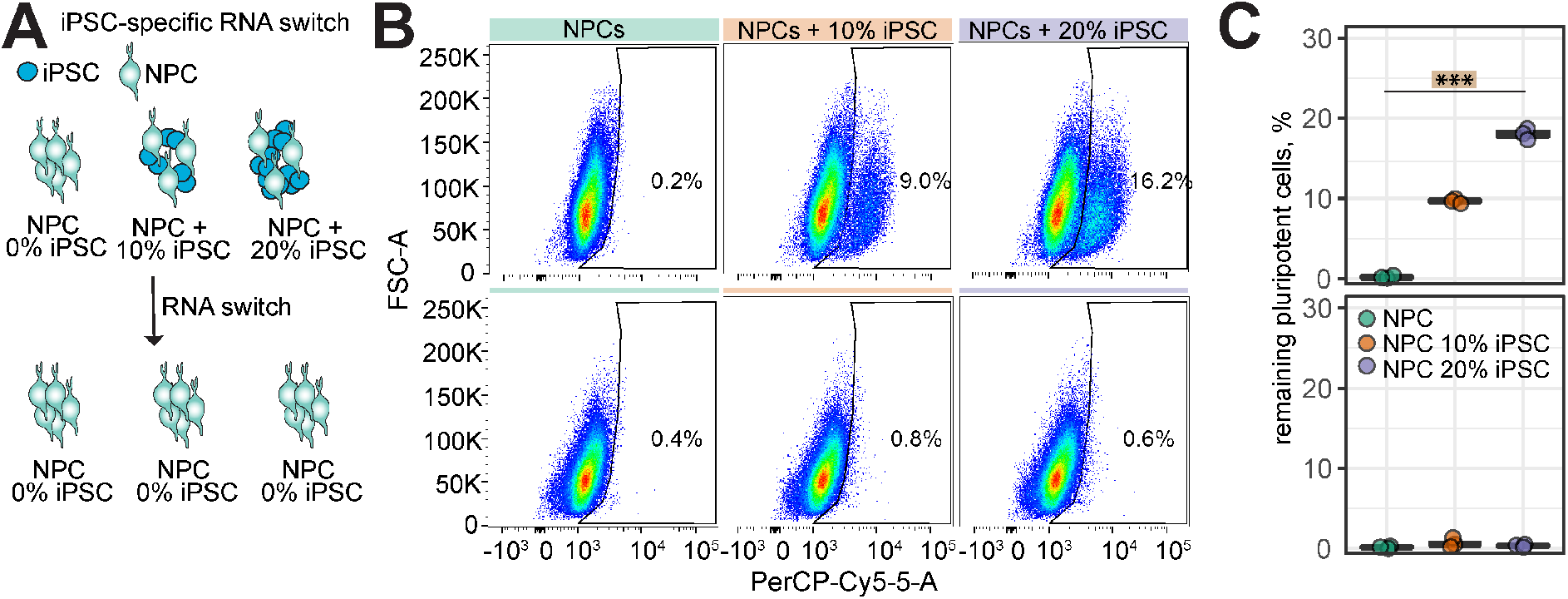
RNA switch for iPSC elimination. A: Schematic representation of RNA switch technology. RNA switch was applied to NPCs without iPSC supplementation (0% iPSC) and with supplementation of 10% and 20% iPSCs. B: Flow Cytometry dot plot of NPCs with negative control switch (upper row) and with RNA switch (lower row). C: Quantification of the remaining percentage of iPSCs in NPC cultures without (top) and with RNA switch (bottom). Boxplots indicate the 25% to 75% quartiles of the data. Each dot in the plots represents one cell culture well and significance of mean differences between the groups was assessed using Tukey’s HSD. Asterisks indicate significance: ** p < 0.01, *** p < 0.001.

### Functionalization of NPCs for in vivo tracking

To be able to track NPCs after transplantation, we established a dual-reporter system combining a fluorescence (GFP) and a bioluminescence (red firefly luciferase (rfLuc)) reporter. We determined the optimal expression conditions and tested luminescence signal intensity for three commonly used promoters, i.e. human phosphoglycerate kinase (hPGK), cytomegalovirus (CMV) and human elongation factor 1 alpha (EF1α) promoter (Fig. 4A). Luminescence signal increased linearly to exposure time, virus titer as well as to the substrate (d-Luciferin) concentration (Fig. 4B, C). The strongest luminescence signals in NPCs were observed when rfLuc was expressed under the EF1α promoter while CMV gave the lowest signal (Fig. 4D). This is in contrast to HEK cells, in which CMV-driven rFluc expression resulted in the strongest luminescence signal, indicating that the promoter has to be selected according to the used cell type. For the following experiments, NPCs transduced with the lentiviral dual reporter under the EF1α promoter were used with a transduction efficiency of ~60-70% (Fig. 4E, F).

**Figure 4:**
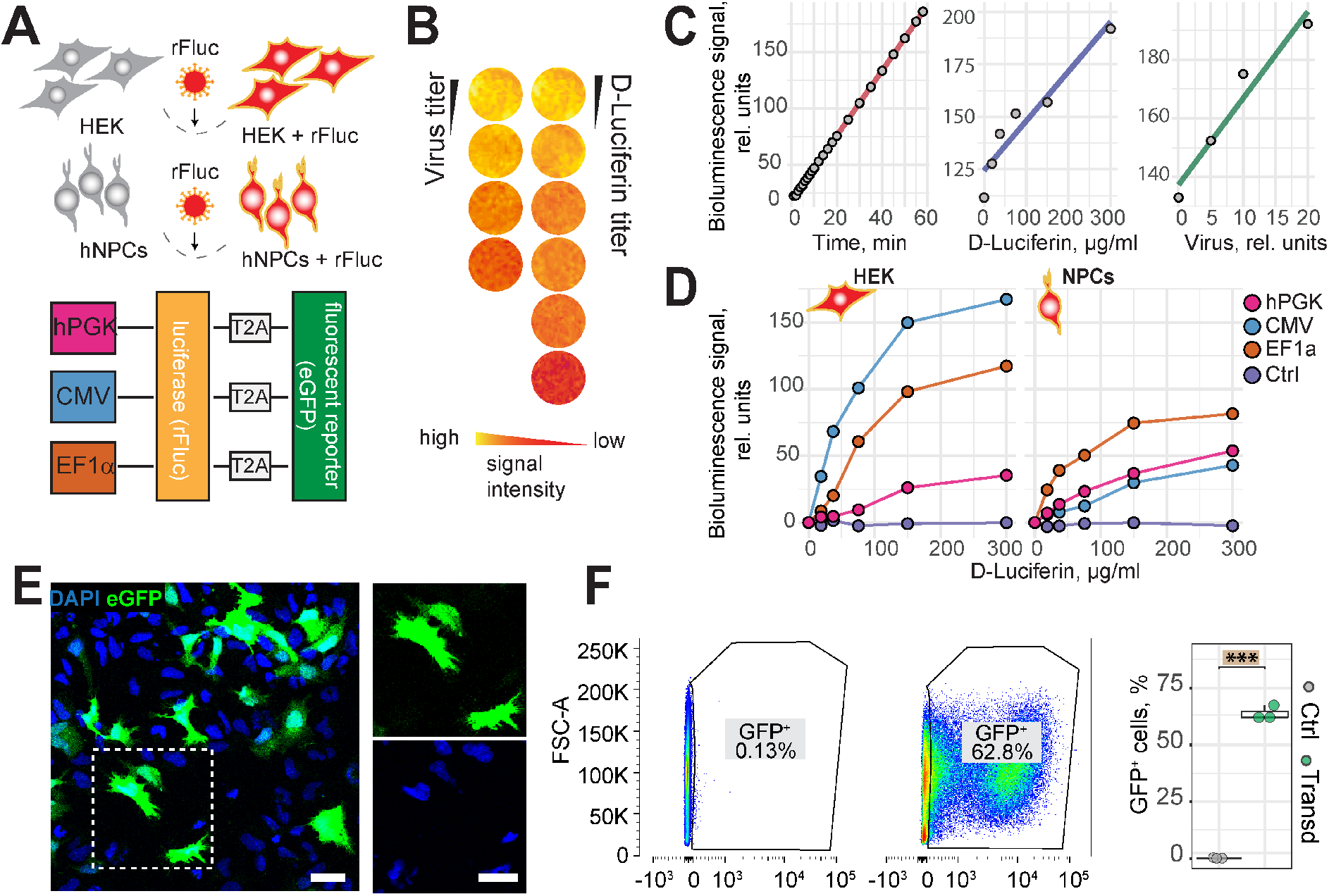
*In vitro* optimization and validation of the dual reporter system. A: Schematic representation of the experimental setup and the lentiviral dual reporter expression constructs. Luciferase and eGFP are expressed under either hPCK, CMV or EF1α promoter. B: *In vitro* luciferase assay, displayed in pseudo-colour. C: Scatter plot including linear regression showing the relation between bioluminescence signal (rel. units) and detection time, virus amount and D-luciferin concentration. D: Luminescence assay showing the luminescence signal in HEK cells (left) and NPCs (right), depending on the used promoter. E: Fluorescence images of transduced NPCs. The rectangle shows an enlarged section as shown on the right. F: Flow cytometry of GFP+ cells in non-transduced (left plot) and transduced cells (right plot). Quantification of GFP+ cells (right graph). Boxplots indicate the 25% to 75% quartiles of the data. Each dot in the plots represents one cell culture well and significance of mean differences between the groups was assessed using Tukey’s HSD. Asterisks indicate significance: *** p < 0.001. rfLuc: red firefly luciferase; hPGK: human phosphoglyceratkinase; CMV: cytomegalovirus; EF1α: elongation factor 1; scale bar: 10 μm; Line graphs are plotted as mean.

### Transplantation and tracking of NPCs in vivo

For in-depth *in vivo* characterization of the NPCs, we transplanted 1.6 × 10^5^ cells stereotactically into the right cortex (AP: + 0.5mm, ML: + 1.5mm, DV: −0.8mm relative to bregma) of wt mice and genetically immunosuppressed NSG or Rag2^-/-^ mice.

As we used freshly thawed cells for transplantation and transplanting various animals can be time-consuming, it was of high importance to make sure that NPC viability was stable for several hours after thawing and before transplantation. Thus, we analyzed the viability of NPCs after thawing over time. NPCs were thawed, kept on ice for 24h and cell viability was measured using the Vi-Cell XR Cell Viability Analyzer at various time points. Viability slightly declined during the first 10h after thawing but remained constant at >75% for further 14h (Suppl. Fig. 2).

To determine the minimum cell number that can be detected by bioluminescence imaging (BLI) *in vivo*, serial dilutions (2 × 10^5^, 1 × 10^5^, 6 × 10^4^, 1.2 × 10^4^, 6 × 10^3^, 2 × 10^3^) of NPCs were transplanted and imaged using bioluminescence imaging. Remarkably, even the lowest cell number transplanted, 2 × 10^3^ NPCs, was detectable (Fig. 5A). A linear relation between bioluminescence signal and transplanted cell number was observed with R^2^=0.98 (Fig. 5B).

**Figure 5:**
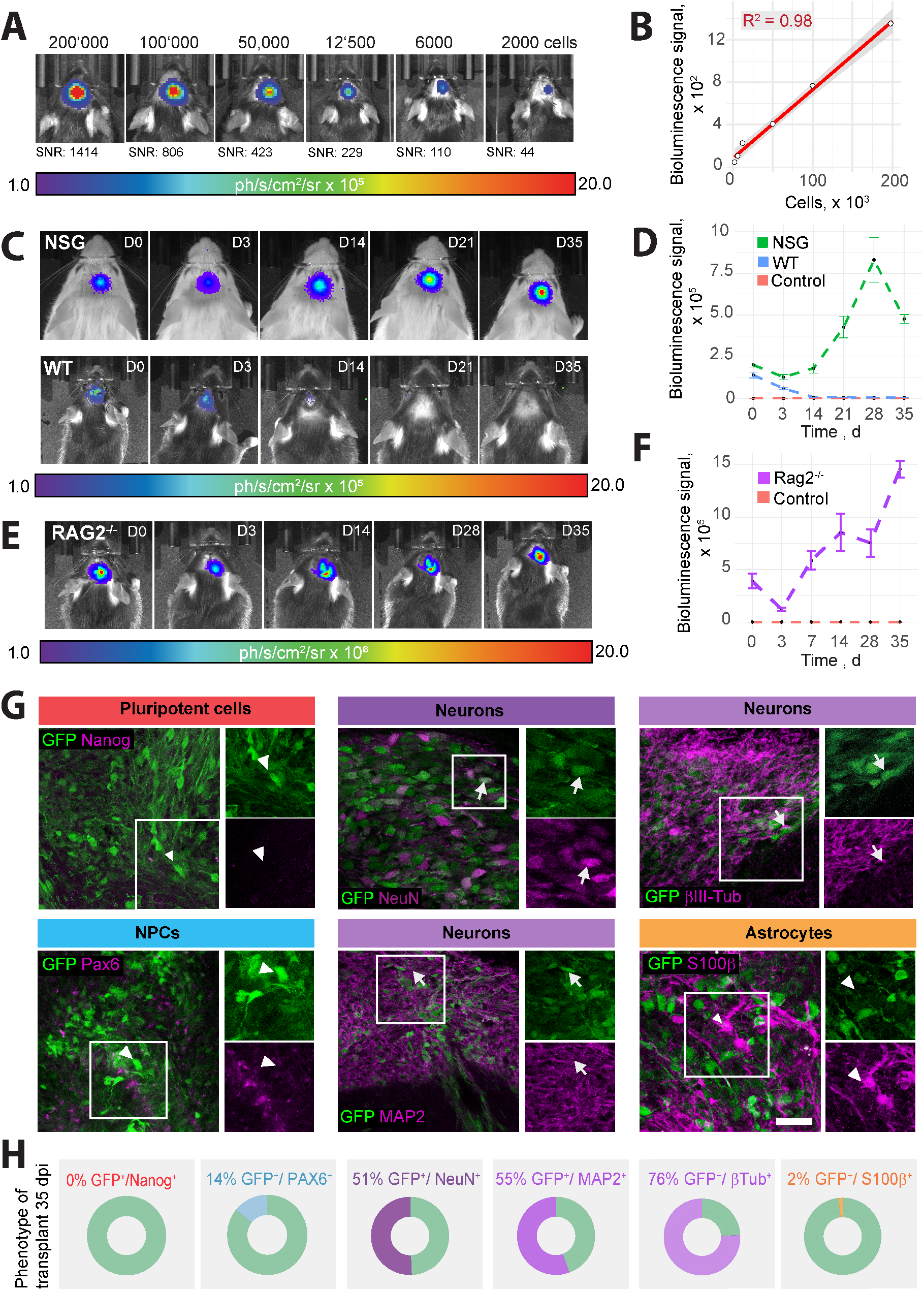
Analysis and *in vivo* tracking of transplanted NPCs. A: Bioluminescence of rFluc-expressing NPCs, transplanted at different numbers into the brain of C57/Bl6 mice at 1.5h post transplantation. B: Scatter plot including linear regression showing the relation between the bioluminescence signal and the number of transplanted cells. C: Bioluminescence of rFluc-expressing NPCs, transplanted into NSG (upper row) and C57/Bl6 mice (lower row), over the course of 35d, measured by *in vivo* bioluminescence imaging. 1.6 × 10^5^ cells have been transplanted. D: BLI signal intensity over time (mean±SEM per strain, SBR). E: Bioluminescence of rFluc-expressing NPCs, transplanted into stroked Rag2-/- mice. F: BLI signal intensity over time (mean±SEM per strain, SBR). G: Representative histological stainings of transplanted NPCs, 35d post transplantation in stroked Rag2^-/-^ mice. Arrowheads indicate GFP-expressing cells that are negative for the respective maker. Arrows indicate positivity for marker and GFP. Scale bar = 50um. H: Quantification of cells positive for the respective marker and for GFP. BLI: bioluminescence imaging; SNR: Signal-to-noise ratio; WT: wild-type; NSG: NOD scid gamma. Line graphs are plotted as mean ± sem.

Survival of dual reporter-expressing NPCs was analysed throughout a 35d time course in wt and NSG mice. While the BLI signal was constantly reduced over time in wt mice and completely decayed between d12 and d21, the BLI signal in NSG mice increased until it reached a plateau between d21 and d35 (Fig. 5C, D). This suggests that immunosuppression is essential for the survival of grafted human NPCs.

To mimic a more clinically relevant scenario, we stereotactically transplanted rFluc-eGFP NPCs in the ischemic brain of stroked Rag2^-/-^ mice. We confirmed a 35-day long-term survival of the graft using bioluminescence imaging, similar to uninjured animals (Fig. 5 E, F).

To assess the long-term differentiation profile of grafted NPCs, we histologically characterized the phenotype of NPCs at 35 days following transplantation in stroked mice. The majority of transplanted GFP^+^ cells expressed various markers of neuronal fate (55% Map2, 76% βIII-Tubulin; 51% NeuN) (Fig 5 G, H). While 14% of transplanted GFP^+^ cells still expressed the neural progenitor marker Pax6, only 2% of cells were positive for astrocyte marker S100β. Importantly, no Nanog^-^-positive cells were detectable at 35 days following transplantation.

In sum, we generated highly pure NPCs from xeno-free iPSCs. These cells can be longitudinally tracked *in vivo* and identified in the brain sections using a dual reporter system. The NPCs have the capability to spontaneously differentiate into functional neurons *in vitro* and *in vivo.*

## Discussion

Despite the high prevalence of neurodegenerative diseases and brain injuries, clinical therapies remain limited. Advances in iPSC technology have brought cell therapy back into the focus of treatment options for the brain [19]. Here, we describe the generation of iPSC-derived NPCs for *in vivo* applications. By modification of previously established monolayer protocols [10,20] we generated highly scalable NPC cultures - yielding 10^18^ NPCs after 15 passages when starting from 10^6^ iPSCs - which are xeno-free and chemically defined. Therefore, this simple monolayer protocol, which does not require cell selection or purification, can smoothly be adapted to GMP-grade for clinical applications. This is of high importance as changes in media composition or in the cell coating during transition from research-grade to a GMP-compliant clinical-grade cell production can strongly affect iPSC differentiation potential and the characteristics of the resulting cells which would complicate the potential clinical translation. We differentiated three iPSC clonal lines to NPCs and showed comparable expression levels of classical NPC marker proteins among all lines. Gene expression analysis revealed a cell type-dependent clustering, clearly distinguishing between iPSCs, NPCs and neural cells while no cell clone-based clustering was observed in NPCs. To determine the potential of NPCs to spontaneously differentiate into neural cells, we withdrew GSK-3β inhibitor CHIR99021, TGF-β inhibitor SB431542, hLIF and FGF2 from the stem cell maintenance medium and cultured the cells for 4 weeks. We found that cells slowed down their proliferation and differentiated into neurons, astrocytes and oligodendrocytes. Expression of calcium sensor GcAMP6 under the human synapsin promoter showed that cells were active and reacted to KCl stimulation indicating the differentiation into functional neuronal cultures. Gene expression analysis further showed a close clustering of differentiated neural cells to human brain lysate, confirming the high translational aspect of iPSC-derived NPCs.

Besides scalability, another crucial point of cell therapy is the purity of the differentiated cells. Residual pluripotent cells in the therapeutic product can cause teratoma formation [19]. The risk for tumours can be strongly reduced by applying highly efficient differentiation and purification protocols. While the differentiation of the iPS cell lines in this study was very effective, evidenced by a strong reduction in pluripotent genes and ≤0.4% Oct4-positive cells in the NPC culture, it cannot be excluded that other iPS cell lines may not yield such a high purity when differentiated to NPCs. Therefore, we tested the compatibility of the obtained NPCs with synthetic RNA switches, a state-of-the-art cell purification technique. The RNA switch technology is a powerful tool for enriching or eliminating a specific cell type in a mixed culture [21,21]. When we added iPSCs to the NPC culture (10% or 20%), the RNA switch resulted in the elimination of iPSCs back to baseline signal. Importantly, when we applied the RNA switch to our NPCs without iPSC addition, we did not observe a difference -flow cytometry still showed 0.2-0.4% Oct4-positive cells. This indicates that the NPC cultures used in this study are already so pure that the RNA switch technique does not have to increase its purity further. Accordingly, we did not observe signs of tumour formation *in vivo*, 4 weeks after transplantation.

An increasing number of studies recently reported the production of NPCs or neural stem/progenitor cells (NSPCs) from either embryonic stem cells or from iPSCs using GMP-compatible techniques [7,22–24]. The first-in-man study using iPSC-derived NSPCs for the treatment of spinal cord injury is currently in preparation at Keio University Hospital, Tokyo, Japan [24]. While many NPC differentiation protocols are laborious and tedious, and require embryoid body formation, neural rosette-selection or colony-picking [7,22–24], the protocol applied here is simply based on a monolayer culture and the treatment with defined neural induction media. A further limitation of several studies, which applied NPSc *in vivo,* is the dependence on endpoint-measures due to the lack of appropriate long-term *in vivo* imaging techniques of graft survival. To circumvent this issue, we used lentiviral-mediated expression of a dual-reporter consisting of rFluc and eGFP, which allowed the *in vivo* tracking of the cell graft via bioluminescence imaging as well as the identification of the grafted cells in brain slices through the eGFP signal. The comparison of different promoters showed that EF1α gave the highest luminescence signal in NPCs while CMV-mediated expression was lowest. Interestingly, this was opposed to our findings in HEK cells indicating that different cell types have different promoter activities. This finding is in accordance with previous observations showing that stem cells can silence exogenous promoters such as CMV [25].

Although we reached a high sensitivity and strong signals during *in vivo* bioluminescence imaging, it was not possible to resolve local cell migration in adjacent brain regions. Furthermore, the bioluminescence signal of other cell types/cell lines may vary due to differences in promoter strength or luciferin metabolism. Detection of cells may also become more challenging in experimental set-ups that require intravenous/systemic injections of cells because of the expected low transmissibility at the blood brain barrier (BBB). These effects may be less pronounced in neurodegeneration or brain injury patients and animal models that are usually associated with BBB damage and increased permeability [15,26].

For our study, we engrafted freshly thawed NPCs, without the need of taking the cells into culture before transplantation. Thus, these cells represent a potential off-the-shelf product for cell transplantation. Cell survival in C57/BL6 mice strongly dropped 14 days post transplantation, while the signal increased in non-injured and in stroked immunodeficient mice. This suggests that the transplanted human cells were attacked by the immune system in non-immunosuppressed mice. It further shows that grafted NPCs survive even in the hostile environment of an ischemic stroke. To exclude that the increased bioluminescent signal in NSG mice is caused by tumour formation, we analyzed histological brain sections. We did not find evidence of tumour formation in histological sections. In contrast, most eGFP-positive, grafted NPCs differentiated into neurons, evidenced by the high positivity for the neuronal markers NeuN, MAP2 and βIII-Tubulin. Therefore, we conclude that the increase in bioluminescence signal is either due to an increased expression in the differentiated cells and/or due to the larger area that neurons cover compared to the smaller NPCs.

Most NPCs, that we transplanted into the stroked mouse brain, differentiated into neurons and only 2% the of cells were positive for astrocyte markers. Several *in vitro* studies have shown that oxygen levels contribute to determining the ratio of neurons and astrocytes, differentiated from NPCs. While 20% oxygen generated a 3:1 ratio of neurons to astrocytes, derived from iPSC-NPCs [27], an increase in the fraction of astrocytes was observed under hypoxic conditions [28,29]. This stands in contrast to the low number of astrocytes generated from the NPCs grafted into the stroked mouse brain in our study. However, reducing oxygen levels in cell culture may not fully mimic the complex environment and the conditions that the graft is exposed to in the stroked brain. Thus, it will be interesting to determine how hypoxic conditions *in vitro* and *in vivo* affect the differentiation potential of our NPCs. Further, follow-up studies are required to determine whether the grafted cells functionally integrate into the neural network and can induce functional recovery in the stroked mice.

## Disclaimers

Hirohide Saito owns shares and is outside director of aceRNA Technologies, Ltd.

## Acknowledgements

We thank Dr. Yoshihiko Fujita, Center for iPS Cell Research and Application (CiRA), Kyoto University, Kyoto, Japan, for providing RNA-switch plasmids. The authors acknowledge funding from Mäxi Foundation, Swiss 3R Competence Center (OC-2020-002) and the Swiss National Science Foundation (CRSK-3_195902).

