## Supplementary data for "Xeno-free induced pluripotent stem cell-derived neural progenitor cells for *in vivo* applications"

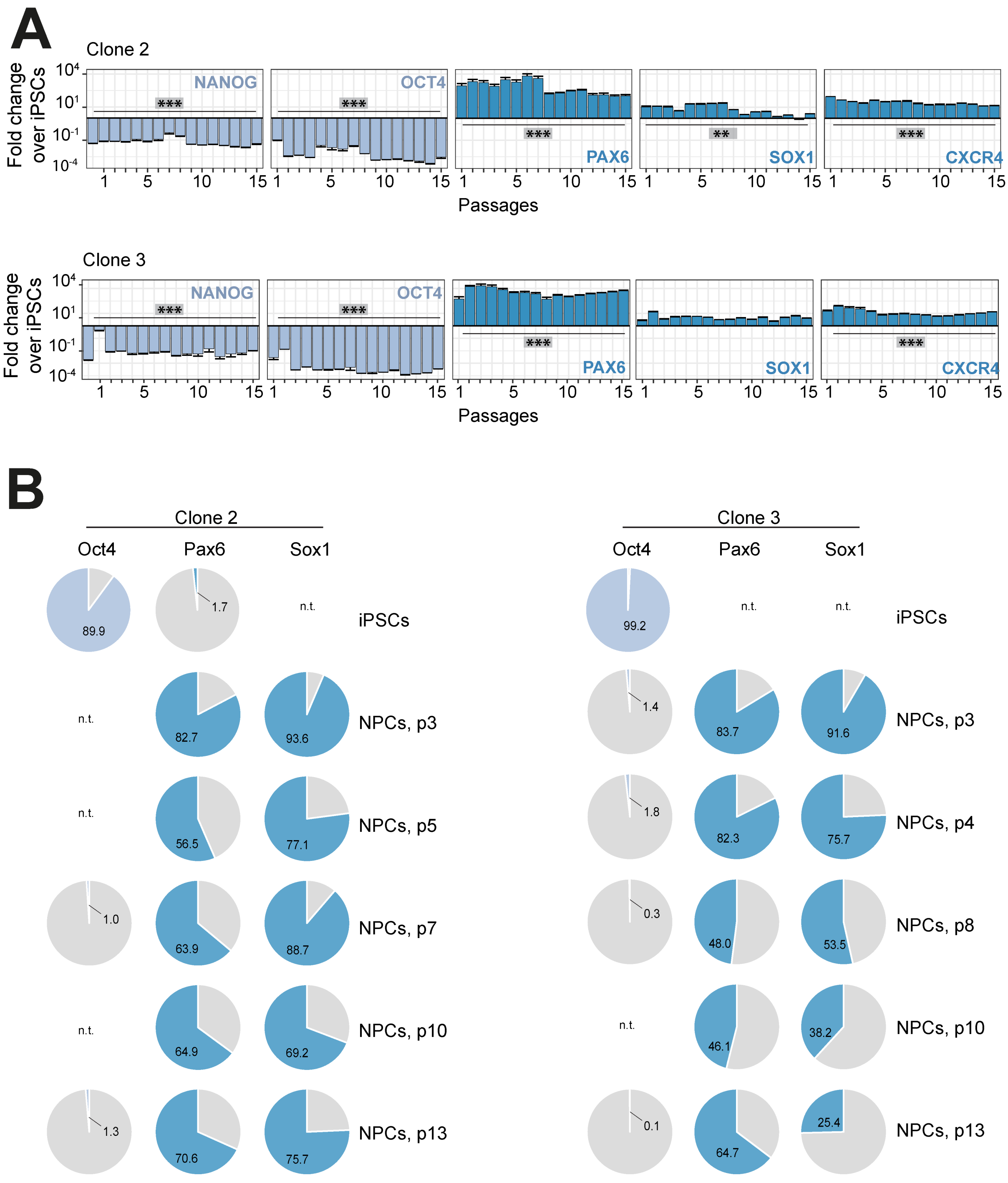


**Suppl. Figure 1: Characterization of iPSC-derived NPCs from two other iPSC clonal lines**
A: Gene expression of pluripotency marker (NANOG and OCT4) and NPC marker (PAX6, SOX1, CXCR4) in NPCs from iPSC clonal lines 2 (upper row) and 3 (lower row) over the course of 15 passages, measured by qPCR. B: Flow cytometry analysis of iPSCs (upper row) and NPCs from iPSC clonal lines 2 and 3 at different passages for Oct4, Pax6 and Sox1. Pie charts illustrate percentage positivity (light/dark blue) for the respective marker and cell type. n.t.: not tested

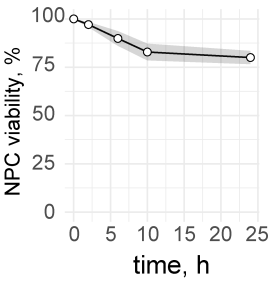


**Suppl. Figure 2: Cell viability *in vitro***. Percentage of freshly thawed viable NPCs over the time course of 25h on ice, measured using Vi-Cell XR Cell Viability Analyzer.

| | |
|---|---|
| DMEM/F12 (50%) 11320-033 | 24ml |
| Neurobasal (50%) 21103-049 | 24ml |
| N2- Supplement (100x) | 500ul |
| B27 – Supplement (50x) | 1ml |
| Glutamax (100x) | 500ul |
| hLIF | 10ng/ml |
| CHIR99021 | 4uM |
| SB431542 | 3uM |
| Dorsomorphin | 2uM |
| Compound E | 0.1uM |

**Suppl. Table 1: Neural Induction Medium 1**

| | |
|---|---|
| DMEM/F12 (50%) 11320-033 | 24ml |
| Neurobasal (50%) 21103-049 | 24ml |
| N2- Supplement (100x) | 500ul |
| B27 – Supplement (50x) | 1ml |
| Glutamax (100x) | 500ul |
| hLif | 10ng/ml |
| CHIR99021 | 4uM |
| SB431542 | 3uM |
| Compound E | 0.1uM |

**Suppl. Table 2: Neural Induction Medium 2**

| | |
|---|---|
| DMEM/F12 (50%) 11320-033 | 24ml |
| Neurobasal (50%) 21103-049 | 24ml |
| N2- Supplement (100x) | 500ul |
| B27 – Supplement (50x) | 1ml |
| Glutamax (100x) | 500ul |
| hLif | 10ng/ml |
| CHIR99021 | 3uM |
| SB431542 | 2uM |
| FGF2 (from passage 2 on) | 5 ng/mL |

**Suppl. Table 3: Neural Stem cell Maintenance Medium (NSMM)**

| | |
|---|---|
| Cell Adhere Buffer | Stemcell Tech |
| Vitronectin XF | Stemcell Tech |
| Accutase Solution | Sigma |
| GlutaMAX-I CTS | Gibco |
| StemMACS™ SB431542 | Miltenyi |
| StemMACS™ CHIR99021 | Miltenyi |
| StemMACS™ Dorsomorphin | Miltenyi |
| N-2 Supplement CTS | Gibco |
| B-27 Supplement XenoFree CTS | Thermo Fisher |
| Animal-Free Recombinant Human LIF | PeproTech |
| StemMACS™ Compound E | Stemcell Tech |
| FGF22 Basic recombinant human protein, Animal-Origin Free | Thermo Fisher |
| Poly-L-ornithine Solution (pLO) | Sigma-Aldrich |
| Laminin-L521 (L-521) | Biolaminin LN |
| Thiazvivin | Sigma-Aldrich |
| DMEM/F12 (50%) | Thermo |
| Neurobasal (50%) | Gibco |

**Suppl. Table 4: Cell culture reagents**
